# Analyzing and Reconciling Colocalization and Transcriptome-wide Association Studies from the Perspective of Inferential Reproducibility

**DOI:** 10.1101/2021.10.29.466468

**Authors:** Abhay Hukku, Matthew G. Sampson, Francesca Luca, Roger Pique-Regi, Xiaoquan Wen

## Abstract

Transcriptome-wide association studies and colocalization analysis are popular computational approaches for integrating genetic association data from molecular and complex traits. They show the unique ability to go beyond variant-level genetic association evidence and implicate critical functional units, e.g., genes, in disease etiology. However, in practice, when the two approaches are applied to the same molecular and complex trait data, the inference results can be markedly different. This paper systematically investigates the inferential reproducibility between the two approaches through theoretical derivation, numerical experiments, and analyses of 4 complex trait GWAS and GTEx eQTL data. We identify two classes of inconsistent inference results. We find that the first class of inconsistent results may suggest an interesting biological phenomenon, i.e., horizontal pleiotropy; thus, the two approaches are truly complementary. The inconsistency in the second class can be understood and effectively reconciled. To this end, we propose a novel approach for locus-level colocalization analysis. We demonstrate that the joint TWAS and locus-level colocalization analysis improves specificity and sensitivity for implicating biological-relevant genes.

## 1 Introduction

With the rapid advancements of sequencing technologies, genetic association analyses have been routinely performed and made significant contributions to gain insights into the roles of genetic variants in complex diseases. Due to recent expansions in large-scale joint genotyping and phenotyping of molecular traits, integrative genetic association analysis has emerged as a tool to study the biological basis of complex diseases. Integrative analyses have the unique ability to interpret genetic associations beyond individual mutations and link complex diseases to functional genomic units, e.g., genes, metabolites, and proteins [1, 2, 3]. Discoveries from integrative genetic analyses have enabled the discovery of novel drug targets [4], hence improved treatments for diseases.

In this paper, our discussions focus on two prevailing types of integrative genetic association analyses: transcriptome-wide association studies (TWAS) and colocalization analysis. Both approaches are widely applied to integrating results from expression quantitative trait loci (eQTL) mapping and complex traits GWAS and show promises in implicating potential causal genes for complex diseases [5, 6]. The analytical goal of TWAS is to test associations between a complex trait of interest and genetically predicted gene expression levels (that are constructed from eQTL information) [7, 8, 9, 10]. More broadly, it connects to the causal inference framework of instrumental variable analysis: given an established TWAS association, the causal effect from a target gene to the complex trait can be estimated [10, 11]. Nevertheless, our discussions focus on the testing stage, which we refer to as *TWAS scan* henceforth. Colocalization analysis aims to identify overlapping causal genetic variants for both molecular (e.g., gene expression) and complex traits [12, 13, 14, 15]. A colocalized genetic variant in a particular cis-gene region implies that a single mutation is responsible for variations in both molecular and complex traits, thus establishing an intuitive link between the traits. A more detailed review of both approaches is provided in the Method section.

Our primary motivation is to investigate the consistency and inconsistency patterns between the inference results by TWAS scan and colocalization analysis in practical settings. Such patterns are examples of inferential reproducibility - one of the three modes defined in the lexicon of reproducibility by Goodman *et al*. [16]. Narrowly speaking, inferential reproducibility refers to the consistency of inference results when different analytical approaches are applied to the same dataset. In practice, both TWAS scan and colocalization analysis are often applied to analyze the same eQTL and GWAS data combinations. However, the implicated genes and the number of discoveries from the two analyses are often markedly different, making biological interpretation and design of follow-up studies challenging. Thus, a systematic investigation to dissect and understand these practical differences is warranted.

Under the settings of inferential reproducibility, the overlapping findings from all approaches are often considered *conceptual replications* for individual methods, with enhanced validity. Nevertheless, the emphasis of the inferential reproducibility analysis is typically placed on the difference sets. We focus on examining the implicated genes reported only by either colocalization or TWAS analysis in our specific application context. Unlike its method and results reproducibility counterparts, inferential reproducibility does not expect or even encourage all methods to yield identical results. On the contrary, the differences driven by different analytical and operational assumptions are largely anticipated [16, 17]. The goal of the analysis is to quantify, understand, and interpret these differences properly. Our study aims to gain insights into how different analytical and operating assumptions of the computational procedures, combined with specific data characteristics, lead to the different implicated genes connecting molecular and complex traits. Specifically, we show that not all the different results in the integrative genetic association analysis are equivalent: in one scenario, they can be reconciled; in the other, they are truly complementary.

Based on our analysis of inferential reproducibility and better facilitate connecting the reconcilable set of genes implicated by the two integrative analysis approaches, we propose a novel *locus-level* colocalization analysis method derived from the same probabilistic modeling framework of fastENLOC. Compared to the existing locus-level colocalization methods in the literature, e.g., RTC [25], JLIM [26], the candidate loci selected for analysis are carefully constructed from the state-of-the-art Bayesian multi-SNP fine-mapping algorithms, and the inference results show much-improved resolution and specificity. Thus, it can serve as a complementary approach to variant-level colocalization, overcoming some of its intrinsic power limitations based on the currently available data [15]. This method is implemented in the software package, fastENLOC (v2.0), and freely available at https://github.com/xqwen/fastenloc/.

## 2 Results

### 2.1 Comparing findings from TWAS scan and colocalization analysis

For this comparative analysis, we select four complex traits, including standing heights from the UK Biobank, coronary artery disease (CAD) status from CARDioGRAM Consortium, and high-density lipoprotein (HDL) and low-density lipoprotein (LDL) measurements from Global Lipids Genetic Consortium (GLGC). These four traits are representative of a range of quantitative and discrete complex traits measured at organismal and molecular levels. We perform both TWAS scan and variant-level colocalization analyses for the selected GWAS traits using the multi-tissue eQTL data from the GTEx project. To enable direct comparisons, we apply the integrative approaches (PTWAS for TWAS scan and fastENLOC for colocalization analysis) utilizing the same set of *cis*-eQTL annotations derived from the multi-SNP fine-mapping analysis method, DAP-G [14].

Our comparison focuses on the gene-level quantification for each trait-tissue-gene combination. The PTWAS scan analysis summarizes the evidence by a *p*-value testing the correlation between a gene’s genetically “predicted” gene expression levels in a specific tissue and the complex trait of interest. In the colocalization analysis, fastENLOC reports a regional colocalization probability (RCP) for each independent eQTL signal of a gene in a specific tissue (also known as a signal cluster) and an independent GWAS hit for a trait of interest. We subsequently compute a gene-level variant colocalization probability (GRCP) by

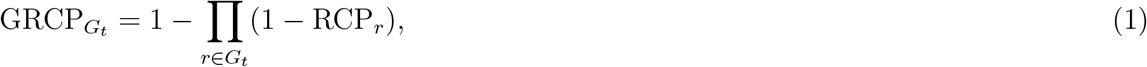

where the set *G_t_* represents the set of all eQTL signal clusters within the cis-region of a gene-tissue pair. The GRCP represents the probability that the gene of interest harbors at least one colocalized causal variant for a trait-tissue-gene combination.

The rank correlations between the −log10 PTWAS *p*-values and the corresponding GRCPs are quite modest, with mean = 0.223 (median = 0.085) across all genes, tissues, and traits. Despite the correlations being significantly different from 0, the low levels of rank consistency seemingly indicate a high degree of discordance (in ranking important genes) between the two types of integrative analyses by the standards of inferential reproducibility.

Next, we inspect the overlapping between noteworthy genes implicated by the two approaches (i.e., the set of conceptually replicated genes). Following [1, 5, 18, 6], we consider that a gene is noteworthy in the colcoalization analysis if its GRCP exceeds the probability threshold of 0.50 for a given trait-tissue pair. In the TWAS analysis, a gene is deemed noteworthy if it is rejected at the FDR 5% level in a trait-tissue pair (FDR controls are performed using the *qvalue* method based on PTWAS *p*-values). The full result of this analysis is summarized in Table 1. Marginally, the TWAS analysis implicates many more noteworthy genes than the colocalization analysis (128,130 vs. 2,337) across 49 × 4 = 196 trait-tissue pairs. Among the two sets, 2054 genes overlap, corresponding to 88% of the noteworthy colocalization genes and 1.6% of the noteworthy TWAS genes, respectively. This finding suggests that most colocalization genes also show strong evidence of TWAS associations, whereas the vast majority of TWAS genes generally lack strong evidence for variant-level colocalizations.

**Table 1:** Noteworthy findings in joint analysis of GTEx eQTL and 4 complex trait GWAS data by different analysis approaches. For TWAS scan, the noteworthy genes are identified at 5% FDR level in each trait-tissue pair; For VC and LC analysis, the noteworthy genes are those with GRCP and GLCP values ≥ 0.50, respectively.

| Complex trait | Analysis approach |  |  |  |  |
| --- | --- | --- | --- | --- | --- |
|  | TWAS | VC* | LC† | TWAS + VC | TWAS + LC |
| Height | 116,396 | 1,674 | 5,387 | 1,524 | 4,701 |
| CAD | 2,500 | 612 | 1,013 | 486 | 762 |
| HDL | 4,996 | 35 | 652 | 30 | 448 |
| LDL | 4,238 | 16 | 464 | 14 | 344 |
| Total | 128,130 | 2,337 | 7,516 | 2,054 | 6,255 |

To better understand the discrepancy between the TWAS and colocalization analysis, we further investigate the two difference sets of the noteworthy genes in greater detail.

#### 2.1.1 Strong colocalization and weak TWAS signals

There are 283 trait-tissue-gene combinations that show strong variant-level colocalization but weak TWAS association evidence. A small subset of these findings can be attributed to the threshold effect of TWAS analysis. That is, if we re-define noteworthy TWAS genes by (slightly) relaxing the FDR control level, these combinations will be re-classified in the overlapping set. However, the majority of the combinations in this set show compatibility with the null hypothesis of the TWAS scan, asserting that genetically predicted gene expression levels are uncorrelated with complex traits of interest. Upon further inspection, we find that most of these instances can be explained by the phenomenon known as “horizontal pleiotropy”.

To illustrate, we take one of the extreme examples in CAD GWAS. There are two independent eQTLs are confidently identified (with posterior probabilities > 0.92) in Gene *TDRKH* (Ensembl id: ENSG00000182134) from the GTEx artery tibial tissue samples. One of the eQTL signals, represented by 10 tightly linked SNPs, also shows strong colocalization evidence, with GRCP = 0.92. In contrast, the predicted gene expression, constructed primarily from these 2 independent eQTL signals, shows little correlation with GWAS CAD status with the resulting *p*-value = 0.98. A detailed instrumental variable (IV) analysis, implemented in the PTWAS estimation procedure, reveals that the two independent eQTLs indicate opposite gene-to-trait effects on the GWAS trait (Figure 1): one implies increased gene expression levels increase CAD risk, whereas the other suggests increased expression levels decrease the risk. When the two eQTLs are combined to predict gene expressions, the overall gene effect on the disease implicated by the predicted expression levels is “canceled out”. The extreme level of heterogeneity in gene-to-trait effects estimated by independent instruments indicates that the vertical pleiotropy represented by variant → gene → trait is highly unlikely in this case.

**Figure 1:**
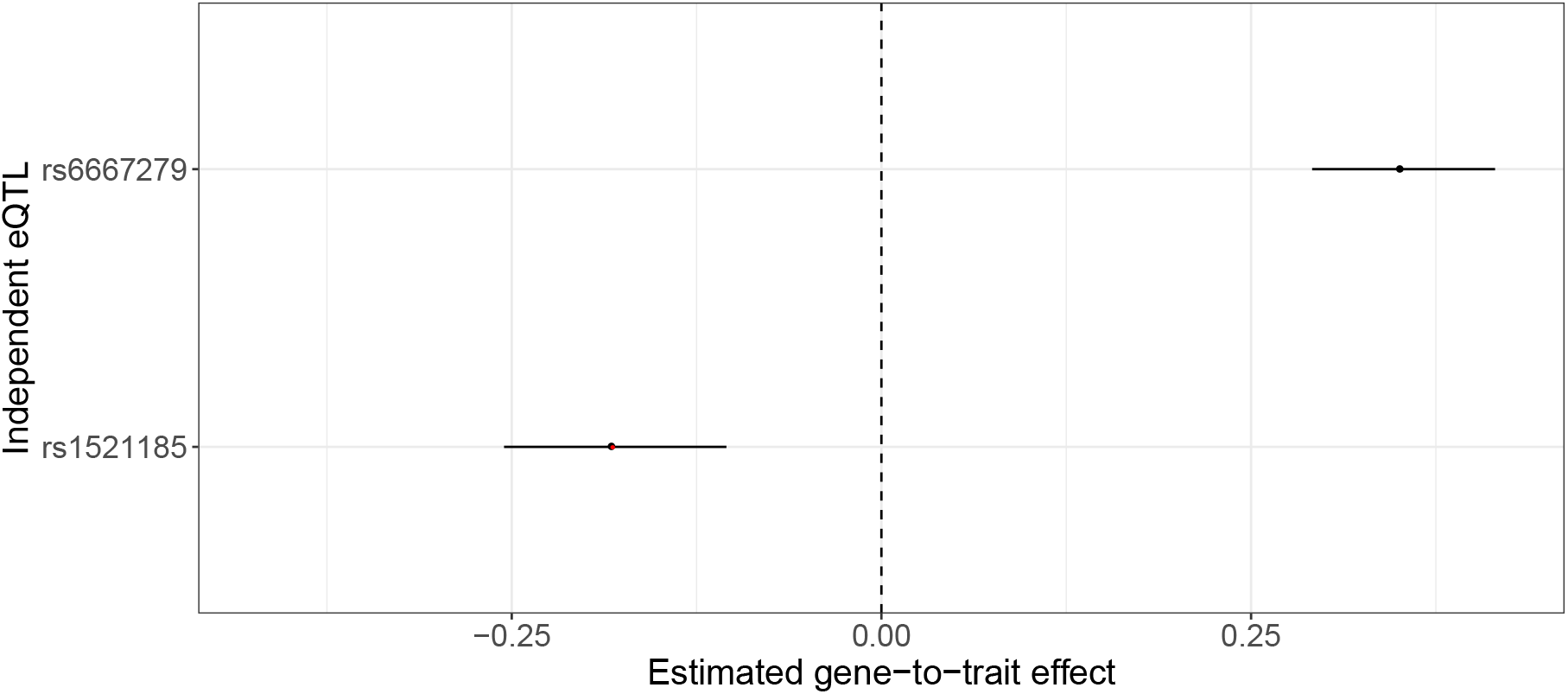
Estimated gene-to-trait effects on CAD by two independent eQTL SNPs of gene *TDRKH* explain its strong colocalization but weak TWAS scan signals. The eQTL fine-mapping analysis of *TDRKH* using GTEx Artery Tibal tissue samples identifies two independent strong eQTLs (SPIPs > 0.9) represented by the lead SNPs rs6667279 and rs1521185, respectively. The second eQTL signal represented by rs1521185 (highlighted in red) also shows strong variant-level colocalization, with RCP = 0.92. The figure shows estimated gene-to-trait effects using the two lead independent eQTL SNPs as instruments. Specifically, the first instrument shows increased gene expression is associated with increased CAD risk, while the colocalized instrument predicts the opposite. As a result, the genetically predicted gene expression by combining the two SNPs shows no evidence of association with CAD risk (PTWAS scan *p*-value = 0.98).

Among the 283 identified combinations, the overwhelming majority of the genes contain multiple independent eQTLs. We quantify the heterogeneity of the inferred gene-to-trait effects using all available independent eQTLs for each combination by computing an *I*^2^ value [10]. (An *I*^2^ value ranges from 0 to 1, where values close to 1 indicate extreme heterogeneity.) For example, the *I*^2^ value for Gene *TDRKH* illustrated above is 0.95. The distribution of *I*^2^ values from this set is shown in Figure 2. The mean *I*^2^ value = 0.73 (median = 0.88), clearly indicating that the majority cases in this category suggest horizontal, instead of vertical, pleiotropy. In comparison, the set of genes implicated by both TWAS scan and colocalization analyses have much lower *I*^2^ values on average, with mean = 0.28 (median = 0.00).

**Figure 2:**
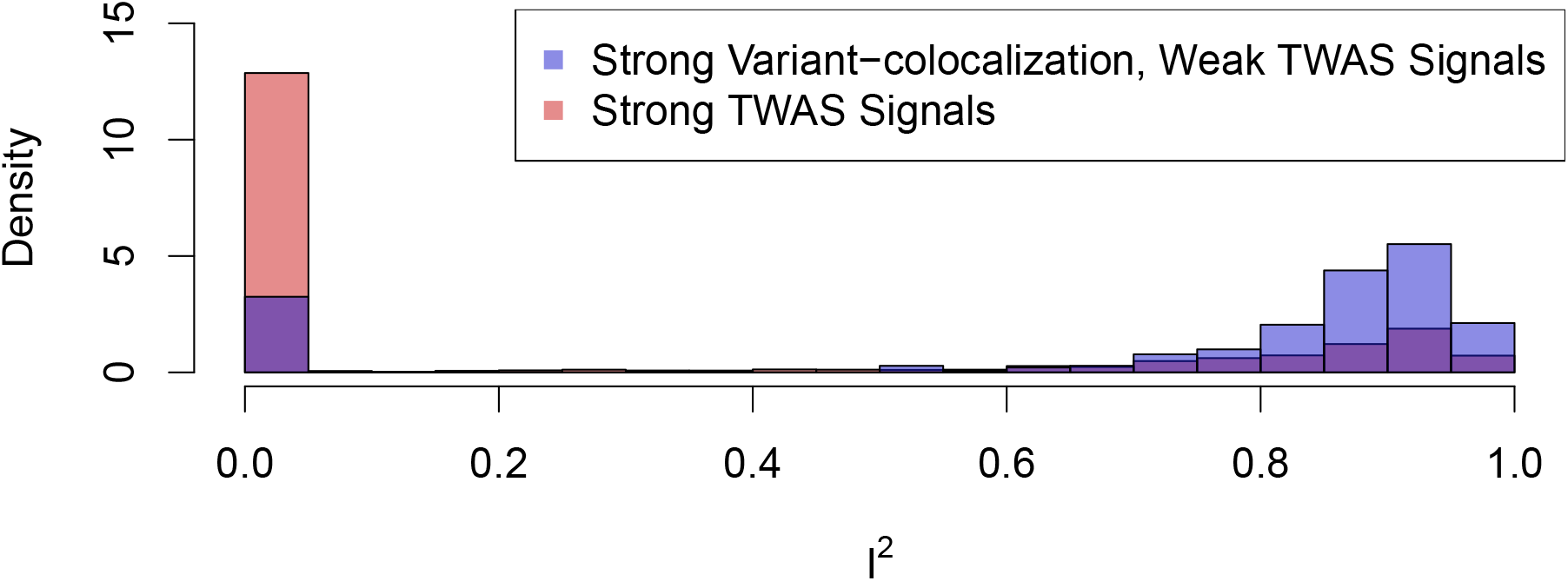
Histogram of *I*^2^ statistics for genes with strong variant-level colocalization but weak TWAS scan signals. *I*^2^ statistic represents the level of heterogeneity in estimated gene-to-trait effects from independent eQTLs. Large *I*^2^ values (i.e., *I*^2^ → 1) indicate that vertical pleiotropy is unlikely. The figure shows that genes with strong variant-level colocalization but weak TWAS scan signals are more likely to have high *I*^2^ values than those with strong TWAS signals. That is, the phenomenon illustrated in Figure 1 can be common in this set of genes. The peak at *I*^2^ = 0 for this set mostly represent the genes whose TWAS scan *p*-value approach but do not exceed the pre-defined FDR significance level.

In summary, we find that most instances in this difference set of implicated genes represent a scenario where the two integrative analysis approaches can be complementary. Additionally, most implicated genes in this set are unlikely direct causal genes to the complex traits, and their relevance to the complex traits could potentially be explained by the phenomenon of horizontal pleiotropy.

#### 2.1.2 Strong TWAS and weak colocalization signals

Strong TWAS and weak colocalization signals account for most discrepancies between colocalization and TWAS analyses. This phenomenon is largely anticipated by different hypotheses employed by the two approaches. A straightforward analytical derivation shows that discovering a TWAS signal does not imply the existence of variant-level colocalization. Instead, the necessary conditions that drive TWAS signals are much relaxed and can be precisely summarized in the following proposition.

##### Proposition 1.

*Assuming linear prediction and association models and provided that a target gene’s genotype-predicted gene expression level is correlated with a complex trait of interest, there is at least one inferred eQTL of the target gene in linkage disequilibrium with a causal GWAS variant*.

*Proof*. Let sets *E* and *G* denote the collections of inferred eQTLs (of the target gene) and the causal GWAS hits, respectively. According to the linear model assumption, the genotype-predicted gene expression can be written as

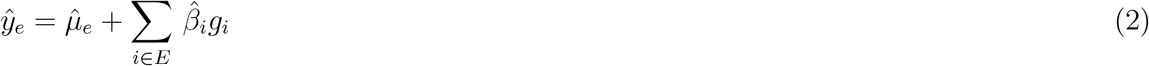

Without loss of generality, assuming the complex trait of interest is quantitative and its genetic association can be described by the following linear model, i.e.,

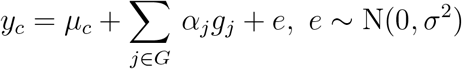

A TWAS scan procedure examines the correlation between 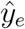 and *y_c_* and tests the null hypothesis, 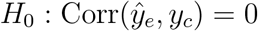. It follows that

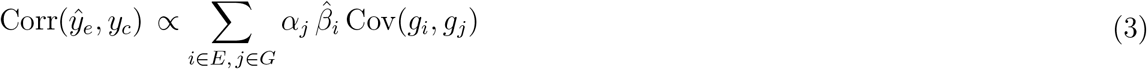

Therefore,

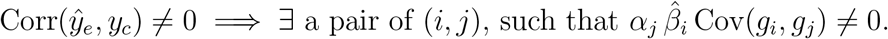

Note that this simple proposition states a *necessary but not sufficient* condition for the existence of TWAS scan signals. Specifically, the TWAS signal is driven by the sum of all non-zero 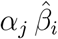 Cov(*g_i_*, *g_j_*) pairs. As illustrated in the previous section, it is statistically possible that multiple terms with different signs can cancel each other out. Additionally, the linearity of the prediction model assumption covers almost all popular TWAS approaches, but it can be relaxed to allow non-linear prediction functions. In the expanded prediction function family, Equation (2) becomes a first-order approximation.

The proposition also implies a direct connection between TWAS scan and variant-level colocalization analysis. By definition, a variant-level colocalization signal satisfies the condition 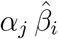 Cov(*g_i_*, *g_j_*) ≠ 0 for some *i* = *j*. A colocalized genetic variant of both molecular and complex traits should also drive a TWAS scan signal in the absence of the cancellation phenomenon. This corollary explains our observation that most genes implicated by the colocalization analysis are also implicated by the TWAS scan.

Next, we consider the scenario that only a single term in (3) drives a TWAS signal by assuming the absence of allelic heterogeneity for the target gene. It becomes apparent that the strength of a TWAS signal reflects the joint effect of *α_j_*, 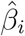, and the LD between the two variants (*i* and *j*). It further implies that even when the LD between a causal eQTL and a GWAS hit is weak, the strength of the resulting TWAS signal can be compensated by relatively strong genetic effects, *α_j_* and/or *β_i_*. This result seemingly explains a common pattern in practical TWAS scan results, where noteworthy signals tend to cluster around some of the strongest GWAS hits. Mancuso *et al*. [19] also discussed a similar pattern of clustered TWAS scan signals due to LD. However, our derivation does not need to assume any true causal relationship between genes and the complex trait of interest (i.e., the phenomenon can exist without any causal genes).

Figure (3) shows a particular instance from the TWAS scan of height GWAS (UKB) and GTEx skeletal muscle genes, where a cluster of TWAS scan signals are centered around one of the most significant GWAS hit on chromosome 3 (rs2871960) identified in the UK Biobank data. The eQTL analysis of the GTEx data suggests the particular SNP is unlikely a causal eQTL.

**Figure 3:**
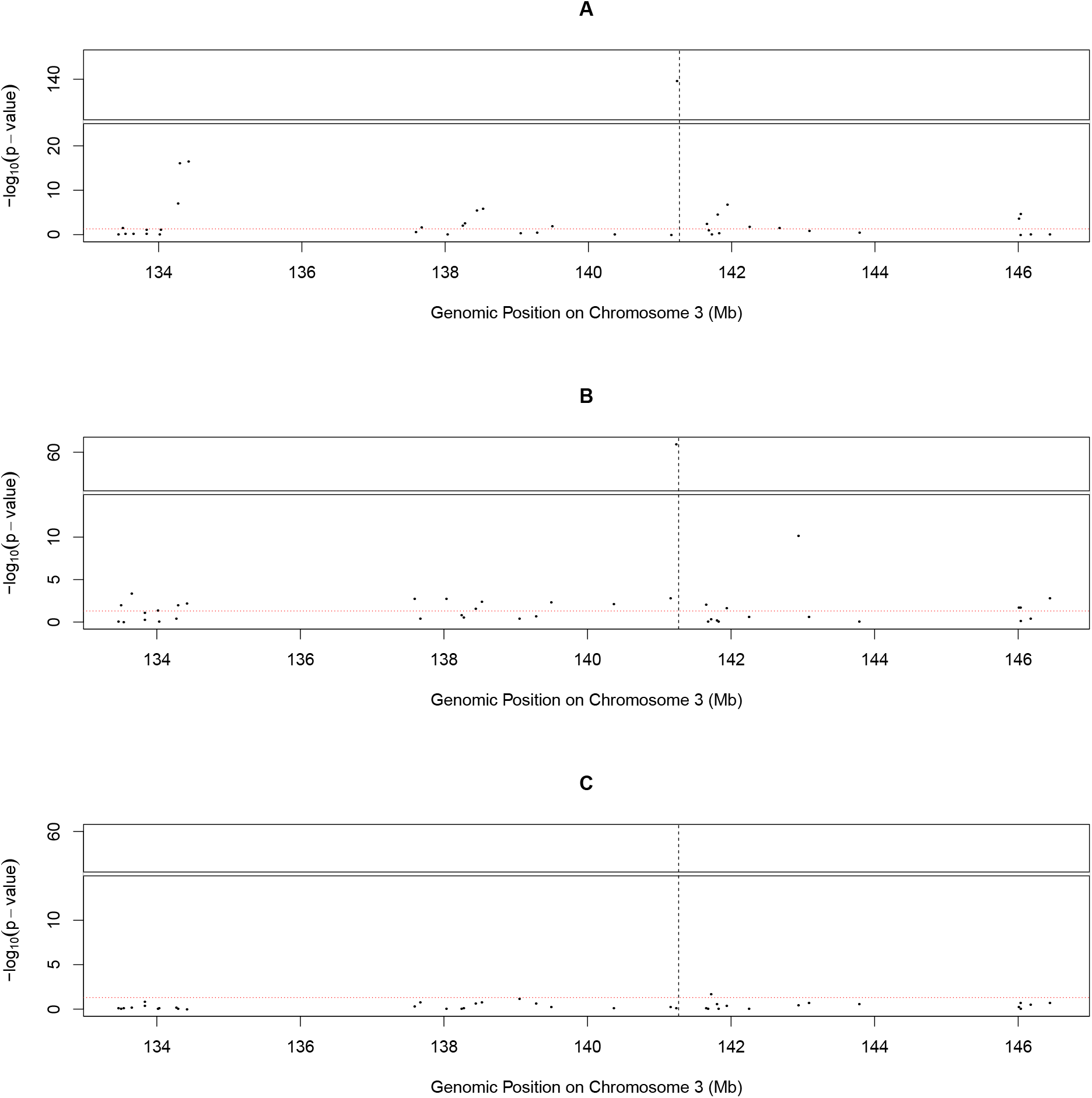
Simulation illustrating LD hitch-hiking effects in TWAS scan. Panel A shows an observed cluster of significant TWAS scan genes from UK Biobank standing height data. The signals are centered around Gene *ZBTB38* (Ensembl ID: ENSG00000177311), whose position is labeled by the dotted vertical line. Each point on the figure represents a TWAS scan *p*-value of a neighboring gene. Panel B shows that a similar cluster pattern can be replicated from a simulated dataset, which is generated by assuming a single causal variant within *ZBTB38*. Panel C shows the significant TWAS scan cluster disappears once the genotypes of the sole causal SNP are regressed out from the simulated phenotype data. In all three panels, the horizontal red line indicates the nominal 0.05 significance level. The simulation experiment illustrates that the significant TWAS scan findings can be attributed to the LD hitchhiking effects.

We conduct simulations to demonstrate that this single GWAS signal (*p*-value = 2.4 × 10^−256^) drives the entire cluster of TWAS signals at this locus. We utilize the real eQTL genotype and phenotype data from GTEx for 39 neighboring genes at this locus and independently simulate the phenotype data of a complex trait with rs2871960 being assumed the only causal SNP. The TWAS scan of the simulated data replicates the pattern observed in the real data, with a third of neighboring genes showing significant TWAS associations. To confirm that the sole GWAS association induces all TWAS signals, we repeat the analysis using the residuals of simulated complex trait phenotype by regressing out the genotypes of SNP rs2871960 (Figure 3).

Although the LD hitch-hiking TWAS signals are not false positives from the perspective of statistical associations, the abundance of signals of these sorts should caution the biological interpretations. For example, it would be mistaken to regard all LD hitch-hiking TWAS signals as *independent* candidates of causal genes for the trait of interest. On this point, we are in full agreement with Mancuso *et al*. [19] that TWAS scan results need to be further processed, better understood, and carefully reported.

In summary, our statistical analysis reveals some main characteristics of TWAS scan signals, which do not require variant-level colocalization and tend to be correlated. These characteristics help explain the difference set of strong TWAS but weak colocalization signals. Regarding this difference set of implicated genes, we find that the two approaches can be reconciled by i) re-defining the standard for colocalization (i.e., adjusting colocalization analysis); and ii) removing non-independent findings from TWAS scan reporting (i.e., adjusting TWAS scan analysis).

### 2.2 Locus-level colocalization: a reconciliation

To reconcile the results between TWAS scan and colocalization analysis, especially targeting the genes showing strong TWAS but weak variant-level colocalization signals, we propose a novel locus-level colocalization analysis method.

From the perspective of variant-level colocalization analysis, the lack of statistical power is a primary limiting factor in analyzing the currently available molecular and complex trait data. Many authors [23, 24] have shown that it is often difficult, if not impossible, to pinpoint the causal genetic associations with the limited sample sizes employed in current molecular QTL mapping studies. Furthermore, with limited sample size, the lead SNPs, whether quantified by Bayesian or frequentist approaches, are often not the true causal SNPs. In many cases, we are relatively certain of a genuine association signal among a group of tightly linked variants (e.g., variants within a signal cluster), however uncertain about the exact variants. Hukku *et al*. [15] show that such uncertainty causes a large class of false-negative findings (i.e., class II FNs) in variant-level colocalization analysis. In their simulation studies based on realistic settings, there are more class II FNs than the identified true findings. Acknowledging this fundamental limitation, we propose the locus-level colcoalization approach aiming to identify the co-existence of casual eQTLs and GWAS hits at an appropriately decreased genomic resolution. The key rationale is that even when locus-level colocalizations do not show strong evidence of variant-level colocalizations in the current data; they may very well be proven class II FNs by future experiments with improved statistical power.

From the perspective of the TWAS scan, the critical issue for reconciliation is to filter out redundant representations due to LD and report only independent and biologically relevant signals. One possible solution is to require causal SNPs for molecular and complex traits colocalized at a small enough genomic region, such that not only is Cov(*g_i_*, *g_j_*) in (3) automatically constrained but the interpretation of the TWAS signal becomes natural. This idea is not new. Many authors [18, 15] have proposed to use variant-level colocalizations as a pre-requisite for following-up TWAS scan results. These proposals are also supported by a class of probabilistic generative models that connect TWAS scan and colocalization analysis (see Eqn (20) in the Method section). Here, we relax the colocalization standard, considering the limited practical power in identifying variant-level colocalizations.

The key to the proposed approach lies in the definition of a genomic locus. As we hope that the variants within a locus represent causal variants for respective traits, a natural solution is to borrow the concept of credible sets/signal clusters from the literature of genetic fine-mapping analysis [20], [14]. The genetic variants classified into a signal cluster are in high LD. More importantly, members of the same signal cluster can be used to interpret the same underlying association signal almost interchangeably with only slight quantitative differences (in model likelihood/posterior probabilities) [20]. By these two properties, at most one variant within a signal cluster represents the true association signal. The candidate loci constructed from signal clusters are also practically small, typically including only a few SNPs in LD. For example, the fine-mapping of the GTEx whole blood eQTL by DAP-G yields 113,318 signal clusters with coverage probability >= 0.95 (i.e., the 95% Bayesian credible sets). On average, each signal cluster contains only 16 SNPs (median = 8) with the minimum pairwise *R*^2^ > 0.5.

The computation of locus-level colocalization probability given pre-defined candidate genomic loci is similar to the analysis of variant-level colocalization. We provide the detailed derivation and description in the Method section. It is worth pointing out that the locus-level colocalization probability (LCP) is fundamentally different from the regional (variant) colocalization probability (RCP). RCP quantifies the probability of a genomic region containing a single colocalized variant. In general, it follows that LCP ≥ RCP for any given loci. For each gene, we define a gene-level colocalization probability (GLCP) for the locus-level analysis, i.e.,

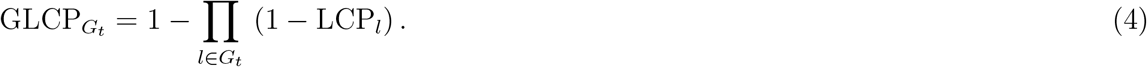

To interpret significant TWAS scan results using the locus-level colocalization analysis, we require that the GLCPs of candidate causal genes exceed a threshold (pre-defined or by FDR control).

### 2.3 Simulation study

We design and conduct simulation experiments to benchmark the proposed locus-level colocalization analysis by comparing it to variant-level colocalization and TWAS scan analyses. We take the real individual-level genotype data from 838 individuals in the GTEx covering 22 distinct LD regions. Specifically, we select random segments of 50 continuous common SNPs (with minor allele frequency > 0.1) from each of the 22 chromosomes. This design aims to ensure modest to strong realistic LD patterns within each LD region but weak LD between different regions. Given the genotype data, we randomly choose 2 causal eQTLs and 2 causal GWAS hits among 1,100 candidate SNPs within each dataset, representing a unique gene. We generate 5,000 datasets where all 4 causal SNPs are located in distinct LD regions (i.e., no variant or locus-level colocalizations). Additionally, we simulate 2,500 datasets with exactly one variant-level colocalization instance, and the remaining causal eQTL and causal GWAS hit reside in different LD regions. Subsequently, the corresponding expression and complex trait phenotypes are simulated by a set of linear models (see details in the Methods section). It is worth noting that this particular simulation scheme can be re-formulated by a structural equation model (SEM) commonly assumed by TWAS scan and Mendelian randomization (Method section). Thus, the simulated datasets are also suitable for TWAS scan analysis. This design is also intended to reduce (or even eliminate) TWAS findings caused by weak LDs between causal eQTLs and GWAS hits.

We analyze the simulated datasets using TWAS scan, variant-, and locus-level colocalization methods. We apply two TWAS scan approaches: PTWAS and the single-SNP TWAS method described in section 1 of the Supplementary Material. For the variant- and locus-level colocalization analyses, we use the algorithms implemented in the software package fastENLOC.

We consider a true positive finding if a simulated gene harboring a colocalized signal is identified at 5% FDR level and a false positive finding if a gene where causal variants reside in distinct LD segments passes the same significance threshold for FDR control. The results by the three different integrative analysis approaches are summarized in Table 2 and Figure 4.

**Figure 4:**
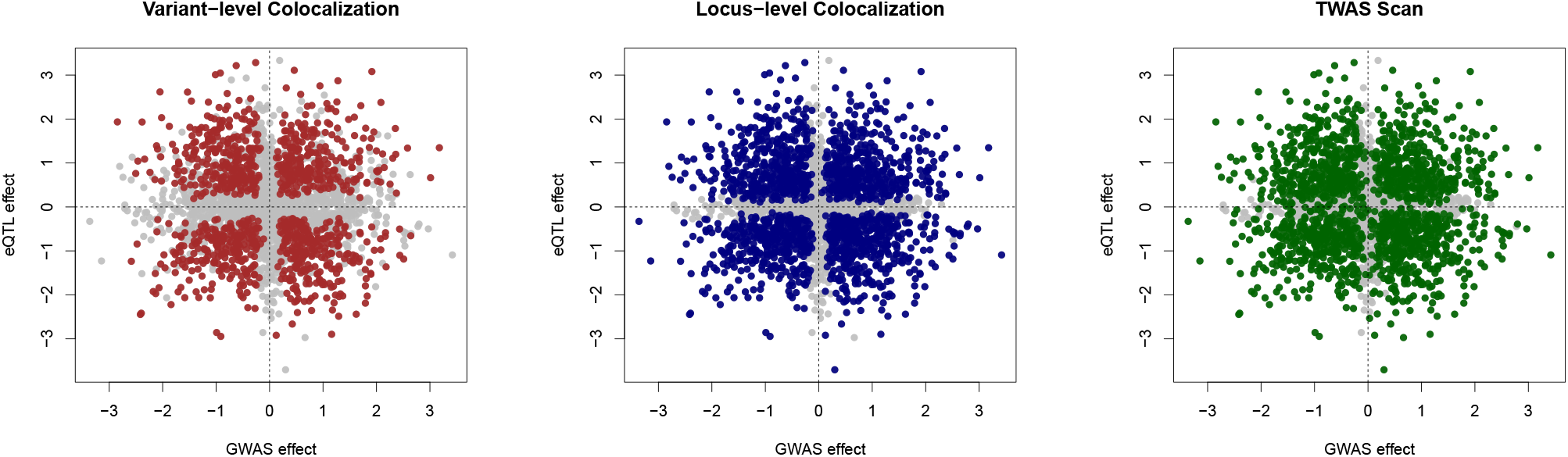
Comparing sensitivity of different integrative analysis approaches in simulations. The bi-plots represent the eQTL and GWAS effects of the colocalized variant in 2,500 simulated datasets. The highlighted points in each plot indicate the true-positive discoveries by the corresponding methods at the 5% FDR level. The power increment of the locus-level colocalization method over the variant-level colocalization method is visually clear. It also exhibits (slightly) improved sensitivity and (much) better specificity over the TWAS scan approach in this simulated context.

**Table 2:**
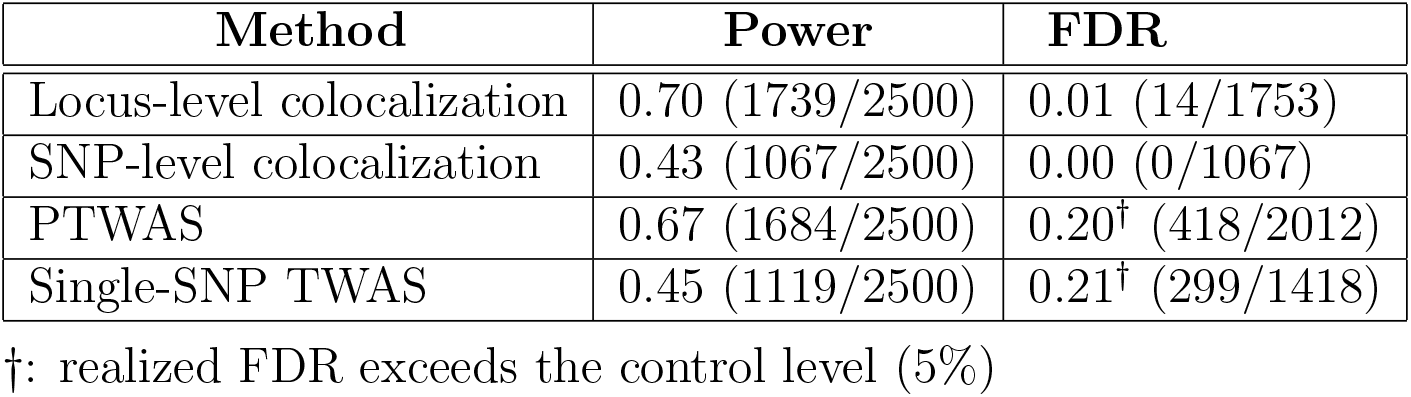
Comparison of power and realized FDR in simulation study.

The variant-level colocalization analysis forms a conservative baseline for comparison. It reports no false-positive findings but the lowest power. Following [15], we further assign all the false-negative findings into two distinct classes. For a colocalized signal, a class I false negative (FN) refers to failure to identify the association signal for at least one trait within a genomic segment; a class II FN refers to failure to quantify the variant-level colocalization probability despite both association signals being localized within the same segment. We estimate that 47.4 % of FNs (or 679 instances) in variant-level colocalization analysis fall into the category of class II. Our results indicate that the proposed locus-level colocalization analysis effectively rescues those class II FNs with little cost in increasing false-positive findings: 95.2 % of the original class II FNs are now identified by the locus-level colocalization analysis. There is no loss of true variant-level colocalization findings, as LCP is always no less than the corresponding RCP (Figure 4).

The multivariate TWAS scan approach reports the most findings (2,102) across all examined methods by a large margin. However, despite our best efforts in assembling the artificial genomic region, a significant proportion of the TWAS scan findings (19.9%) are results of weak LDs between distinct eQTL and GWAS variants located in different segments. Specifically, we find that the maximum *R*^2^ values between a causal eQTL and a causal GWAS hit range from 6 × 10^−4^ to 0.148 (mean = 0.027) within this set. A detailed inspection confirms that the true expectations of gene expression levels (computed by the true genetic association model of gene expressions) are indeed significantly correlated with the simulated phenotypes. We emphasize that these findings are only considered false positives by the standard of colocalization analysis or the *intended* structural equation model. In summary, the extremely low level of LD required to drive a TWAS scan signal can be surprising, but this phenomenon is well explained and anticipated by Proposition 1 (i.e., the strong genetic effects of eQTLs and/or GWAS hits can compensate weak LDs). Interpreting the (biological) relevance of this set of genes can be difficult, as such associations should only be characterized as accidental. The joint analysis by filtering TWAS scan results using locus-level colocalization analysis results is proven effective in removing these accidental association signals. The filtered TWAS discovery set maintains reasonably good power (61%) and is much easier to interpret.

### 2.4 Re-analysis of GTEx and GWAS data using locus-level colocalizations

Finally, we re-analyze the 4 complex traits GWAS and the GTEx eQTL data using the proposed locus-level colocalization approach. We again ensure that all examined methods utilize identical input information. Detailed comparisons between PTWAS, variant-, and locus-level colocalization analyses are provided in Table 1. Across 196 trait-tissue pairs, the locus-level colocalization identifies 7,516 genes with GLCP > 0.50, representing a 2.2 fold increase of discoveries than the variant-level colocalization analysis. A remarkable 83% of the high GLCP genes overlap with the significant PTWAS scan genes. We consider this set of 6,255 PTWAS genes filtered by locus-level colocalization analysis as high-priority genes for further validation.

We first inspect the set of 1,261 genes that show high GLCP but do not pass FDR control at 5% level in PTWAS scan analysis. Similar to the weak TWAS and strong colocalization signals implicated by variant-level colocalization analysis, the is set of genes also shows excessive heterogeneity in estimated gene-to-trait effects by multiple independent eQTL signals (Figure 5), suggesting potential horizontal pleiotropy. Figure 5 also indicates an increase of genes whose PTWAS scan signals are close but do not pass the FDR control threshold.

**Figure 5:**
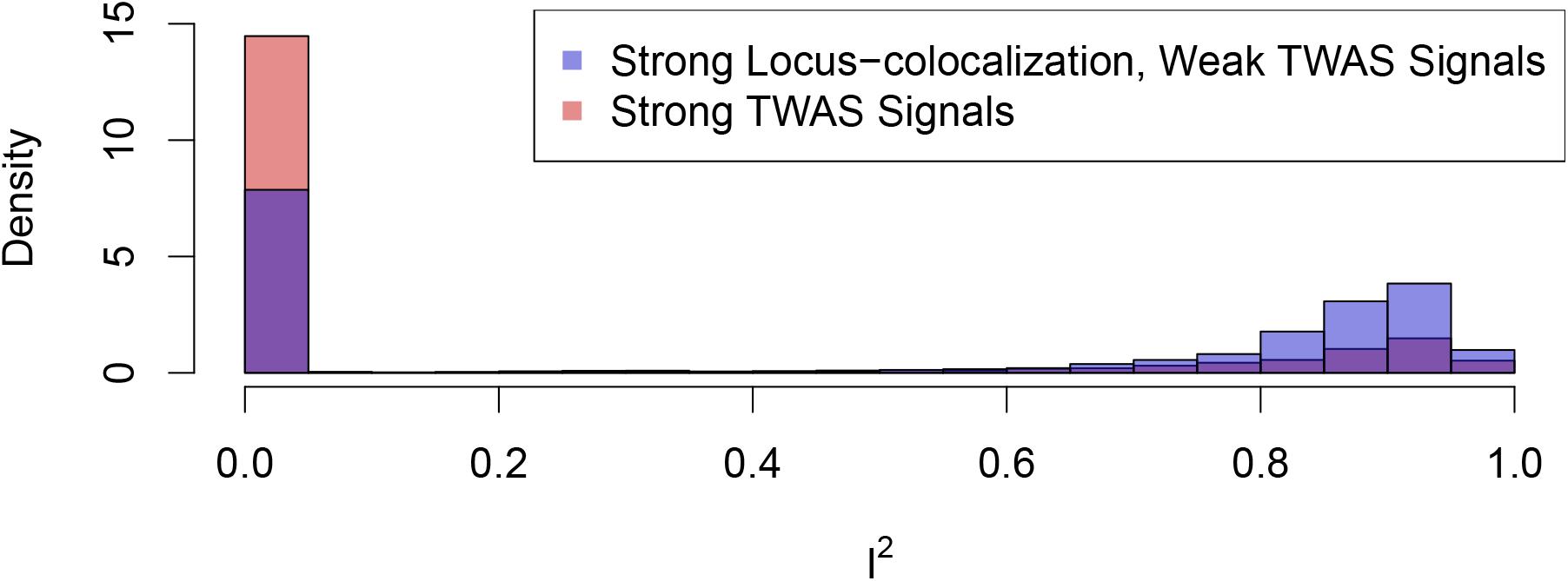
Histogram of *I*^2^ statistics for genes with strong locus-level colocalization but weak TWAS scan signals. The figure indicates that genes with strong locus-level colocalization but weak TWAS scan signals are more likely to have high *I*^2^ values than those with strong TWAS signals. This phenomenon is similar to the genes with strong variant-level colocalization but weak TWAS scan signals, indicating both sets of genes are likely enriched with cases of horizontal pleiotropy.

In the set of 6,255 filtered PTWAS significant genes by the locus-level colocalization analysis, we find many new discoveries have documented biological relevance to the corresponding complex traits in the literature. We first investigate the Online Mendelian Inheritance in Man (OMIM) [21] database to find associated genes. For this investigation, we select phenotype OMIM IDs mapped to any of our four traits and subsequently identify all *confirmed* gene associations with those phenotypes. In total, we extract 12 validated genes from the OMIM across the 4 analyzed complex traits, with 6 in our result by the joint PTWAS and locus-level colocalization analysis (Supplementary Table S1). It is worth noting that the joint SNP-level colocalization and PTWAS analysis only identifies 1 out of 12 genes.

An illustrative example is gene *LPL* (Ensembl id: ENSG00000175445) with HDL. Within the adipose subcutaneous tissue, there is a very strong TWAS signal (*p*-value = 9.7 × 10^−135^) and substantial evidence for a locus-level colocalization (GLCP = 0.97). Meanwhile, the GRCP for this gene is only 0.07, reinforcing the added utility of locus-level colocalization.

Additionally, we utilize one of the largest gene-disease association repositories, DisGeNET [22], to inspect the biological relevance of CAD-associated genes implicated by the proposed joint-analysis. DisGeNET comprehensively integrates and ranks multiple types of reliable gene-disease association evidence from a catalog of source databases. We pull out a list of 65 high-confident CAD-relevant genes, whose DisGenNET scores are all greater than the built-in default selection threshold of 0.3. In our integrative analyses, the PTWAS scan identifies 510 unique genes across 49 tissues. A subset of 172 unique genes passes the additional filtering of the locus-level colocalization analysis. We find that 23 of the 172 genes appear in the DisGenNET CAD gene list (Supplementary Table S2). In comparison, the remaining 338 genes that lack locus-level colocalization evidence only contribute 7 additional hits on the DisGenNET CAD list. A Fisher’s exact test indicates that the enrichment of CAD-relevant genes implicated by the proposed joint-analysis is statistically highly significant in contrast to stand-alone TWAS scan analysis (*p*-value = 8.8 × 10^−7^). Compared to the joint-analysis of TWAS and variant-level colocalization analysis (which flags 17 DisGenNET genes from 117 implicated unique genes), the level of signal enrichment is similar, but the confirmed discoveries by the newly proposed method represent a 35% increase.

## 3 Discussion

This paper systematically investigates two prevailing types of integrative genetic analysis methods, TWAS scan and colocalization analysis, focusing on understanding and reconciling their inferential differences in practical settings. From the perspective of inferential reproducibility, we identify multiple statistical and biological factors that yield different sets of implicated genes. In one scenario (i.e., strong colocalization but weak TWAS signals), we find that most genes in the specific difference set show interesting characteristics requiring further biological investigations, indicating the two approaches are complementary. In the other scenario (i.e., strong TWAS but weak colocalization signals), we find that the differences can be effectively reconciled. Subsequently, we propose and implement a new locus-level colocalization analysis method to bridge the two types of analyses. We illustrate that the proposed joint-analysis approach can produce a rich list of biologically relevant “conceptual replications” for downstream investigations and validations.

Variant-level colocalization analysis utilizes a conceptually rigorous and superior standard to examine the overlapping of causal association signals at the finest resolution. It exhibits the highest specificity among the existing integrative analysis approaches. The most noticeable drawback is its limited sensitivity given currently available data [15]. The issue is originated from the difficulty in quantifying variant-level association evidence/uncertainty in the presence of LD [23, 24]. While future data with more precise phenotyping (e.g., single-cell expression data) and/or larger sample size will certainly improve the power of variant-level colocalization analysis, the intrinsic difficulty due to complex LD patterns may not be fully resolved. We trade off the rigid conceptual standard of variant-level colocalizations for improved sensitivity in the proposed locus-level colocalization analysis. This is mostly motivated by a series of realistic simulation studies presented in [15] and this paper, where the ratios of class II false negatives versus reported findings (which contain few false-positive errors) are often strikingly high (i.e., ~ 1).

We acknowledge that locus-level colocalization analyses are not new. For example, RTC [25] and JLIM [26] are two representatives of general-sense locus-level colocalization methods in the literature. However, we see at least two distinct advantages of the proposed approach over the existing methods. First, we define target loci through the fine-mapping output of signal clusters/credible sets, which harbor very limited but highly relevant candidate genetic variants. In comparison, existing approaches often analyze genomic regions at hundred kb scales [25, 26, 9]. As expected, the precise locus definition by the proposed approach results in high specificity, which is much comparable to variant-level colocalization analysis as shown in our real data analysis. Second, the proposed approach is based on the same Bayesian hierarchical model that explicitly estimates the enrichment of molecular eQTLs in GWAS hits and subsequently utilizes the enrichment information to boost the discovery of colocalized sites. This is a unique and novel feature of the proposed locus-level colocalization method.

The empirical comparison of TWAS scan and colocalization analyses helps us identify statistical factors that differentiate the two sets of results. One of this work’s most important take-away messages is that TWAS scan results are unlikely independent and need further processing. At a certain level, PTWAS scan is analogous to the practice of single-variant testing in the common practice of genetic association analysis, where the community standard is *not* stopping at reporting significant individual variants but summarizing the testing results and grouping linked variants to flag independent causal variant-harboring loci.

Our proposal to apply locus-level colocalization to screen and filter TWAS scan results follows the similar strategy established in PhenomeXcan [18], which is proven effective in identifying biologically relevant potential causal genes. Given the increased sensitivity of locus-level versus variant-level colocalization analysis, the improvement in the performance of such a strategy is logically expected. We note that the software FOCUS utilizes an alternative and statistically elegant strategy (analogous to the multi-variant fine-mapping in genetic association analysis) to parse correlated TWAS scan findings and identify potential “causal” genes. This strategy can be highly effective if the assumption of a causal gene in the genomic region of interest is met. In comparison, our proposed strategy does not require such an assumption but relies on the biological implication from colocalizations, hence offering some added robustness in inference. In practice, the two strategies can be complementary and applied simultaneously. Additionally, we note that many authors [8, 9, 10, 11] have connected TWAS analysis to Mendelian randomization (MR) and instrumental variable (IV) analyses and point out the scan procedure is equivalent to the *testing* procedure in MR and IV analyses. We strongly agree that additional estimation and heterogeneity diagnosis procedures from the MR and IV analyses can be further applied to validate the causality of implicated genes in combination with the proposed joint-analysis procedure. We believe that the conceptual replications identified from the proposed joint-analysis forms a good set for such purpose. Finally, joint TWAS scan and colocalization analysis can be used to identify cases of horizontal pleiotropy by examining genes with strong colocalization but weak TWAS signals. Such findings can be critical to uncover the full molecular mechanisms underlying complex diseases and deserve attention from the community.

Much of this work is motivated by understanding the inferential reproducibility between TWAS scan and colocalization analysis. Unlike the other types of reproducibility, namely, methods and results reproducibility, inferential reproducibility does not prompt pursuing the consistency of outcomes from different analytical methods [16]. Instead, the primary aim is to identify analytical assumptions/factors driving inferential differences and understand the extent of inconsistency between methods. These factors are particularly important for practitioners to design analysis schemes with all available tools. As we have demonstrated in this paper, inferential reproducibility is fundamentally different from inferential errors/mistakes. It should be treated with care, and most importantly, in a context-dependent manner.

## 4 Methods

### 4.1 Overview of TWAS scan and variant-level colocalization analysis

This section provides a brief overview of TWAS scan and variant-level colocalization analyses. There are multiple implementations for each integrative approach. Here, we emphasize the commonality among different implementations and refer the readers to the cited publications for their differences.

#### 4.1.1 TWAS scan analysis

TWAS scan analysis aims to identify genes whose genetically predicted expression levels are associated with a complex trait studied in a GWAS. Most available approaches assume a linear prediction model for gene expression levels. Consider a target gene with *m cis*-SNPs. The general form of a linear prediction function using *cis*-SNP genotypes is given by

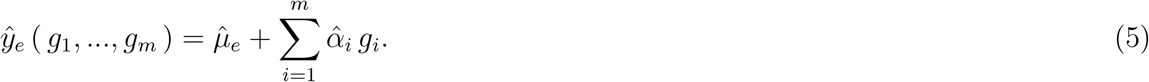

This supervised learning model is subsequently trained using the available eQTL datasets. Different TWAS scan approaches apply different learning algorithms to train the prediction model and estimate the parameters, 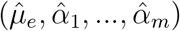. For example, PrediXcan utilizes the shrinkage method, elastic net; TWAS-Fusion, PTWAS adopt Bayesian prediction and model averaging approaches, respectively. The fully trained prediction model can be applied to GWAS datasets and impute the gene expressions using only the corresponding genotype information. Finally, association testing is performed between the observed complex trait phenotype and the imputed expression phenotypes. Particularly, under the linear prediction model assumption, the association testing procedure can be effectively carried out using only summary-level GWAS statistics instead of explicit imputation and testing using individual-level genotype-phenotype data from GWAS.

Existing literature has explicitly connected TWAS scan to the testing procedure in instrumental variable analysis/Mendelian randomization [8, 10, 11]. Particularly, the prediction model (5) is viewed as a composite instrumental variable (i.e., a weighted sum of genotypes) and is shown more powerful than the methods considering single genetic variants as eligible instruments. Conversely, a prediction model using a single genetic variant can be straightforwardly derived following the principles of both supervised learning and instrumental variable analysis (Section 1 of the Supplementary Material). Because this approach is computationally trivial and theoretically more powerful than the existing SMR method, we use this approach to represent the single-variant TWAS scan method in this paper.

Finally, we emphasize that the TWAS scan does not represent the full IV/MR analysis procedure, where gene-to-trait effect estimation and model checking is typically required. Nevertheless, under the high-dimensional setting of integrative genetic association analysis, these additional procedures can be subsequently applied on TWAS scan results [10].

#### 4.1.2 Variant-level colocalization analysis

Variant-level colocalization analysis aims to identify the overlapping of causal eQTL and GWAS SNPs. The problem is challenging due to wide-spread linkage disequilibrium and the limited power in uncovering true causal associations given the currently available molecular and complex trait data. Most existing variant-level colocalization methods take the Bayesian probabilistic modeling approach to effectively account for the inevitable uncertainty in genetic association analysis [14, 15, 12, 13]. They also take advantage of fine-mapping results obtained from individual analysis of each trait to achieve improved accuracy. The colocalization probabilities for individual SNPs (i.e., SNP-level colocalization probabilities or SCPs) can be unimpressive, especially when a few SNPs are in high LD. The aforementioned methods all report a regional-level colocalization probability (RCP) to represent the probability that a genomic region harbors a single colocalized variant.

Variant-level colcoalization analysis is known to have limited power given the currently available GWAS and molecular QTL data. There are two classes of false-negative (FN) errors commonly encountered in practice [15]. Specifically, the class I FNs refer to the cases where the genetic association analyses for individual traits fail to identify at least one genuine association at a colocalized site. The class II FNs are caused by inaccurate quantification of association evidence at the individual variant level: even both associations are identified for a locus, the probabilistic characterization of the assumed causal variants may lead to strong evidence against variant-level colocalization.

#### 4.1.3 Evaluating inferential reproducibility between TWAS scan and colocalization analysis

In practice, both TWAS scan and colocalization analysis are applied to the same data, which forms the basis for evaluating their inferential reproducibility. In this paper, we select the TWAS scan method, PTWAS scan [10], and the variant-level colocalization method, fastENLOC [18, 15], for this purpose. In our real data analysis, both PTWAS and fastENLOC utilize the same probabilistic eQTL annotations derived from the GTEx project (v8) [1] and the fine-mapping method, DAP-G [27]. The GWAS summary statistics for the 4 selected complex traits are publicly available and further harmonized to match variants genotyped in the eQTL dataset. Our main motivation for these selections is to minimize the procedural differences in eQTL and GWAS data pre-processing. Thus, we can focus on the important analytical factors that lead to differences in inferential results.

### 4.2 Probabilistic colocalization analysis at locus level

In this section, we outline the analytical derivation and computational algorithm for evaluating locus-level colocalizations.

#### 4.2.1 Defining genomic loci

Our implementation defines a locus by the intersected signal clusters from the target molecular and complex traits. A signal cluster is defined by a set of LD SNPs representing the same underlying association signal for a given trait. Intuitively, any member of a signal cluster can similarly represent an independent association signal in likelihood computation, but the true causal SNP may not be distinguishable. This construction of loci for colocalization analysis implies that each resulting locus harbors at most one causal SNP for the complex trait and at most one causal molecular QTL.

#### 4.2.2 Computation of locus-level colocalization probability

We denote the genotype and phenotype data combinations for the molecular and the complex traits of interest by ***Y**_e_* and ***Y**_g_*, respectively. Consider *p* overlapping SNPs in a predefined locus. Let *β_i_* and *α_i_* denote the genetic effects of SNP *i* for the complex and molecular traits, respectively. The latent binary association status of all member SNPs in the locus for the complex trait is represented by a *p*-binary vector, *γ*, where *γ_i_* = **1**(*β_i_* = 0). Similarly, we use *p*-vector ***d*** to denote the latent association status for the molecular trait, where *d_i_* = **1**(*α_i_* = 0).

Let Γ_*k*_ and *D_k_* denote the sets of configurations of *γ* and ***d*** values with exactly *k* independent association signals for the corresponding traits, respectively. By the construction of the genetic locus through fine-mapped signal clusters, it follows that

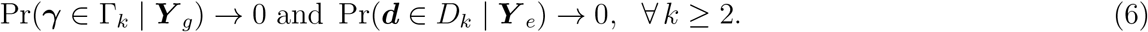

The problem of locus-level colocalization then can be framed as evaluating the following posterior probability, Pr(*γ* ∈ Γ_1_, ***d*** ∈ *D*_1_ | ***Y**_g_*, ***Y**_e_*). Assuming the molecular and complex trait data are collected from two non-overlapping sets of samples, it follows that

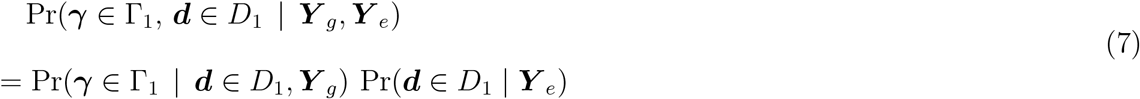

Next, we show that both required probabilities can be deduced from the variant-level colocalization model detailed in [14, 15].

It follows from the locus definition and (6) that

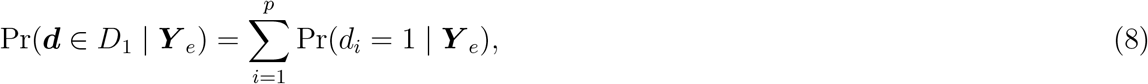

where Pr(*d_i_* = 1 | ***Y**_e_*) denotes the posterior inclusion probability (PIP) for member SNP *i* computed from the fine-mapping analysis of the molecular QTLs.

The computation of Pr(*γ* ∈ Γ_1_ | *d* ∈ *D*_1_, ***Y**_g_*) can be formulated as a problem of fine-mapping GWAS hits with informative molecular QTL priors [27]. More specifically, the variant-level priors,

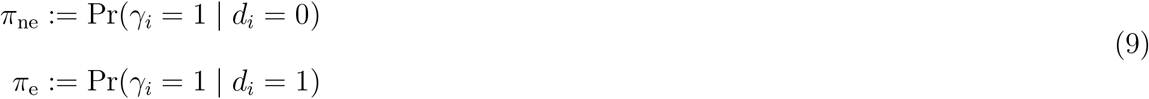

can be estimated by a multiple imputation (MI) procedure described in [14]. Thus, the prior required for the locus-level colocalization analysis is computed by considering all compatible configurations of *γ* and ***d*** values, i.e.,

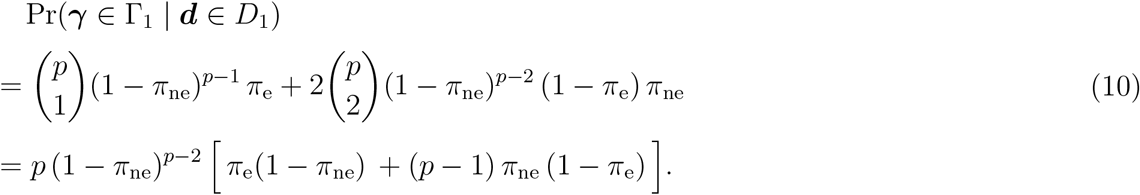

Similarly,

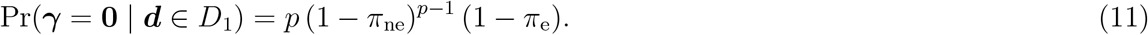

As in the variant-level colocalization analysis, the PIPs of GWAS SNPs are assumed available from Bayesian fine-mapping methods based on an exchangeable and eQTL non-informative prior, Pr(*γ_i_* = 1):= *π*, which can also be estimated by the average of all GWAS PIPs [14]. For this eQTL non-informative prior, the induced prior probabilities for the locus of interest are given by

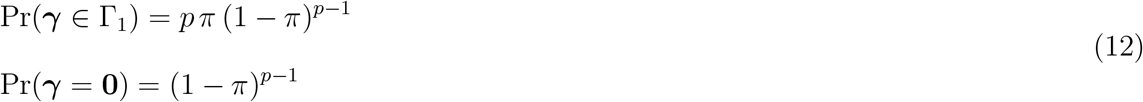

It again follows from the locus definition and (6) that

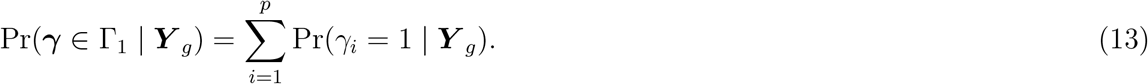

Consequently, the marginal likelihood defined by the following Bayes factor,

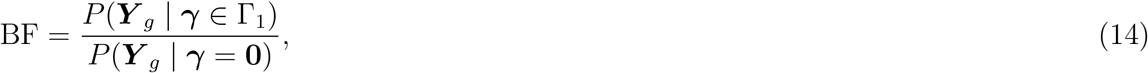

can be obtained by

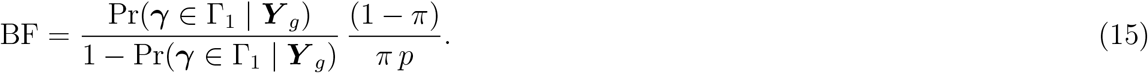

Put together, by (6), (10), (11), (15), and the Bayes theorem,

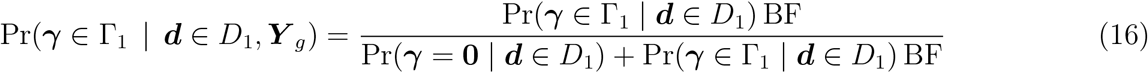

Consequently, the desired locus-level colocalization probability, Pr(*γ* ∈ Γ_1_, ***d*** ∈ *D*_1_ | ***Y**_g_*, ***Y**_e_*), can be computed by (7).

### 4.3 Simulation details

#### 4.3.1 Simulation to illustrate LD hitchhiking effect in TWAS scan

Our simulation scheme is informed by real data analysis. Firstly, we identify the SNP rs2871960 as one of the most significant genetic associations (*z* = −34.2) for standing height in the UK Biobank data. In the GTEx data, this SNP maps to the cis-region of a single gene, *ZBTB38* (Ensembl ID: ENSG00000177311), in the Muscle Skeletal tissue. Additionally, we identify 38 neighboring genes, all within 8 Mb of the SNP. A list of all 39 genes is provided in the Supplementary Table S3.

In this simulation, we consider all 22,662 *cis*-SNPs of the 39 genes. The eQTL dataset is directly taken from the GTEx muscle skeletal tissue with 706 individuals. We note that rs2871960 is unlikely to be the true causal eQTL for ENSG00000177311 in the muscle skeletal tissue based on the fine-mapping result. Specifically, it does not fall into any signal cluster of the gene and the PIP = 2.75 × 10^−3^ (despite its single-SNP testing *p*-value reaching 10^−11^). Additionally, the SNP-level colocalization probability for this SNP is also quite low (2.72 × 10^−3^). We simulate a complex trait (***y**_h_*) for the 706 GTEx individuals using their true genotypes (***g***) for SNP rs2871960 and the following simple linear regression model,

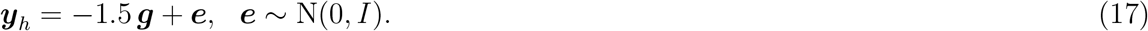

That is, the only causal SNP for the complex trait is assumed to be rs2871960, with its genetic effect fixed at –1.5. Note that the genotypes for the gene expression and complex trait are perfectly matched. However, because the complex trait is independently simulated, the overall scheme fits the two-sample design for integrative analysis. The particular genetic effect value attempts to match the observed complex trait *z*-score in the UK Biobank with much-reduced sample size. To further illustrate that all significant TWAS findings from the simulated dataset are due to the LD hitchhiking effect, we regress out the genotypes of the causal SNP and treat the residuals as a new complex trait phenotype. Finally, we analyze both datasets by PTWAS scan and report the corresponding *p*-values for each examined gene.

#### 4.3.2 Simulation to evaluate locus-level colocalization analysis

In this simulation, we assemble a genomic region with a “known” LD structure from the genotype data of 838 GTEx samples. We classify the artificial genomic region into 22 LD blocks, with each block containing 50 consecutive common SNPs from a unique chromosome. We intend to capture natural LD patterns within each block and minimize LD between blocks (as the genotypes are taken from distinct chromosomes). The LD structure of the assembled genomic region is shown in Supplementary Figures S1 and S2.

For each simulated dataset, we independently generate phenotype data for a molecular (***y**_e_*) and a complex trait (***y**_c_*) using the following linear regression models

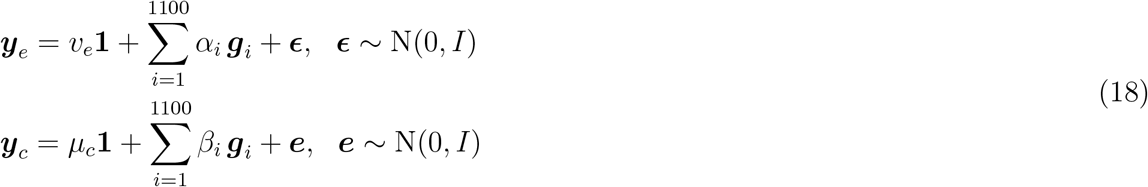

For each trait, only 2 SNPs located in different LD blocks have non-zero genetic effects. The exact effects for the two causal SNPs of each trait are independently sampled from the distribution N(0,1). As a result, the percentage of variance explained (PVE) by genetics for each trait is approximately 0.35 on average. Coincidentally, we find that under this signal-to-noise ratio, the GLCP threshold at the 5% level FDR control is very close (only slightly higher) to the commonly used value of 0.50 in colocalization analysis. We simulate 5,000 datasets with the causal eQTLs and the GWAS hits located in distinct LD blocks (i.e., no colocalization). They serve as a baseline to examine potential false-positive findings. For the other 2,500 datasets, we explicitly select one causal eQTL SNP and one GWAS SNP colocalized at a single variant while the remaining casual eQTL SNP and GWAS SNP are placed in distinct LD blocks. Let ***g***_1_ denote the colocalized SNP, the generative models for the phenotypes become

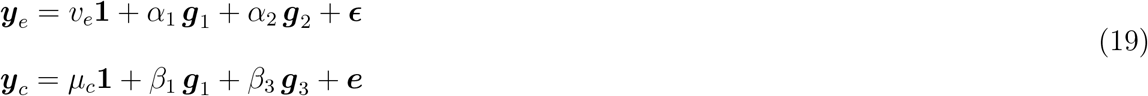

Note that the colocalization in (19) induces a structural (mean) equation between ***y**_e_* and ***y**_c_*, i.e.,

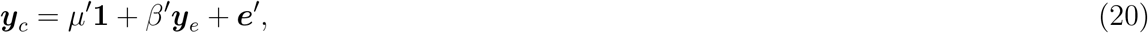

which is often used in TWAS simulations [8, 9].

### 4.4 Analysis of GTEx eQTL and complex trait GWAS data

We use the multi-tissue eQTL data generated from the GTEx project (v8) in our real data analysis. The data are processed and analyzed by the GTEx consortium. The pre-processing and analysis protocols are documented in [1]. For evaluating inferential reproducibility, we particularly focus on the cis-eQTL fine-mapping results generated by the software package DAP [27], which are publicly available at the GTEx portal. These results are re-formatted to feed PTWAS scan and fastENLOC without further processing.

The summary statistics from the 4 selected complex trait GWAS (HDL and LDL from the GLGC consortium; standing height from the UK Biobank; coronary artery disease from the CARDioGRAM consortium) are also publicly available. We use the single SNP association *z*-scores for the 4 traits harmonized by the GTEx project [1]. The main purpose of the harmonization procedure is to match common SNPs between the SNPs interrogated in the GWAS and the GTEx project. More specifically, if a GTEx SNP is missing from the corresponding GWAS, its GWAS *z*-score is imputed using the software package impG. The harmonized *z*-scores are subsequently used in PTWAS scan and fastENLOC analyses.

