## Supplementary Material for "Analyzing and Reconciling Colocalization and Transcriptome-wide Association Studies from the Perspective of Inferential Reproducibility"

### Supplemental Material

### 1 Single-variant TWAS scan

In this section, we provide details for the single-variant TWAS scan procedure used in the paper. It serves as a representative baseline for all TWAS scan approaches. The simplicity of the procedure is particularly appealing and helps elucidate some of the key common features of TWAS scan analysis.

We view the described procedure under the special constraint that only a single variant is allowed for gene expression prediction. Under this setting, the resulting TWAS scan $p$-value is simply the single-SNP GWAS association $p$-value of the most significant eQTL SNP identified from the single-SNP eQTL mapping.

We note that the best single-SNP gene expression prediction model is represented by the most significant eQTL SNP based on the training eQTL data. This is because the single-SNP association $p$-vallue is monotonic to the $R^2$ value (or PVE) of the corresponding simple regression model, which directly measures the model's predictive ability. We denote the resulting optimal single-SNP prediction model by

$$\hat{\boldsymbol{y}}_e = \hat{\beta}_e \, \boldsymbol{g}_e, \tag{1}$$

assuming all gene expression phenotypes are pre-centered.

In a separate GWAS sample, TWAS scan measures the correlation between phenotype vector, $\boldsymbol{y}_c$ and the vector of the predicted gene expression, $\hat{\boldsymbol{y}}_e = \hat{\beta}_e \boldsymbol{g}'_e$. Because $\hat{\beta}_e$ is a constant, it follows that

$$\text{Corr}(\boldsymbol{y}_c, \hat{\boldsymbol{y}}_e) = \text{Corr}(\boldsymbol{y}_c, \boldsymbol{g}'_e). \tag{2}$$

This very quantity and the corresponding standard error can be equivalently measured by a simple regression model, regressing $\boldsymbol{y}_c$ on the selected eQTL SNP genotypes.

Alternatively, the procedure can be explained by an optimal single instrument Mendelian randomization (MR) testing procedure. The MR testing procedure examines the null hypothesis that the specified genetic instrument is uncorrelated with the complex trait of interest (i.e., the outcome) [1, 2]. It should be clear that the most significant eQTL SNP is the optimal (or the strongest) if candidate instruments must be single genetic variants (i.e., no composite instrument allowed). This reasoning also leads to the same procedure.

##### 1.1 Comparison to SMR

Summary data-based Mendelian Randomization (SMR) also utilizes a single genetic variant for testing. Thus, it falls into the category of single-SNP TWAS procedure. It is also derived from the principle of MR but instead relies on the MR estimation procedure to derive the test statistic. The resulting test statistic derived by [3] is given by:

$$z_{gy}^2 \frac{z_{gx}^2}{z_{gy}^2 + z_{gx}^2} \sim \chi_1^2 \tag{3}$$

under the null.

It is clear that the test statistic from the optimal single-variant TWAS approach follows that

$$z_{gy}^2 \sim \chi_1^2, \tag{4}$$

under the MR null hypothesis.

However, because

$$z_{gy}^2 \frac{z_{gx}^2}{z_{gy}^2 + z_{gx}^2} \leq z_{gy}^2 \qquad \forall z_{gx}, z_{gy}, \tag{5}$$

it implies that the described procedure is universally more powerful. This result should not be surprising, as our derivation has also indicated the optimality of the proposed procedure.

It should be noted that the SMR offers additional inference features, i.e., a gene-to-trait effect estimate and its corresponding standard error. Nevertheless, because our focus is on the TWAS scan/testing procedure, the proposed single-variant TWAS approach offers much simpler computation and results in a more powerful test.

### Supplementary Figures

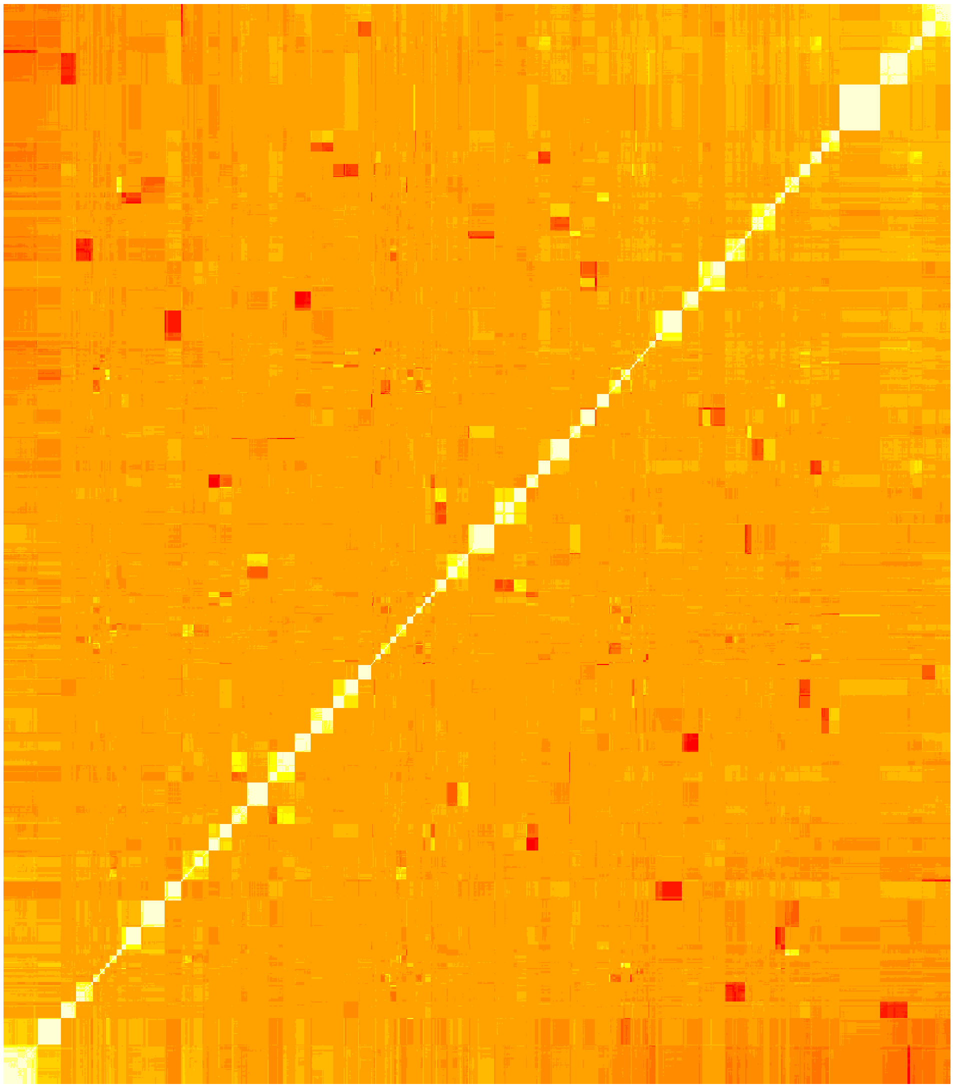

Figure S1: **LD structure of assembled genome used in the simulation study - overall view** LD is measured by pair-wise $r^2$ between SNPs. The assembled genome contains 22 LD blocks. Each block is selected from a different chromosome and consists of 50 consecutive common SNPs (MAF $\geq 0.1$). The genotype data for the selected SNPs are taken directly from 838 GTEx muscle skeletal samples.

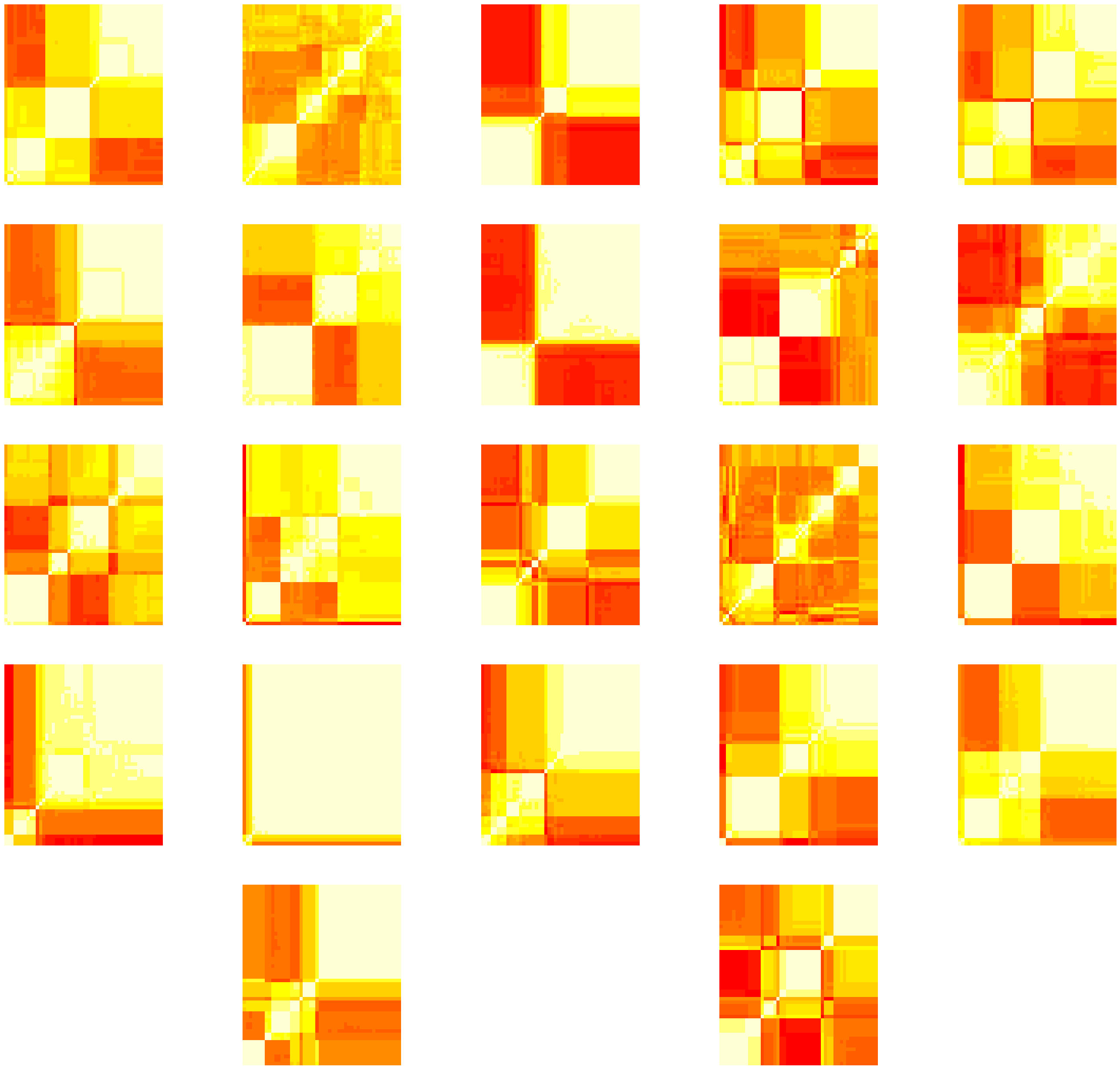

Figure S2: **LD structure of assembled genome used in the simulation study - breakdown by LD blocks** Each figure in the panel shows LD structure within an LD block, consisting of 50 SNPs, in the assembled genome.
